# Appraising the natural root-knot nematode resistance in *Solanum sisymbriifolium*, a wild relative of potato

**DOI:** 10.1101/2024.06.20.599809

**Authors:** Itsuhiro Ko, Allan B. Caplan, Joseph C. Kuhl, Cynthia Gleason

## Abstract

Root-knot nematodes (RKNs) are a major pest of Solanum and other economically important crops worldwide. Two species of RKNs (*Meloidogyne chitwoodi* and *Meloidogyne hapla*) are persistent threats to potato growers of the United States. These RKNs infect potato roots and tubers, causing tuber blemishes that decrease potato market value and significantly impact the profitability of the infected potato crop. Due to environmental, health, and economic concerns, the longstanding control methods of using soil fumigants and post-plant nematicides are not favored by producers and consumers. Therefore, deploying RKN resistant cultivars is an alternative method to control RKN damage. However, there is no genetic resistance to RKN in commercially-available, cultivated potatoes. Therefore, the critical first step to breed a RKN resistant plant is to identify a genetic source of RKN resistance. A wild *Solanum* species, *Solanum sisymbriifolium,* also known as litchi tomato, can effectively control several agronomically important species of plant parasitic nematodes. *Solanum sisymbriifolium* is completely resistant to RKNs; only a few nematodes enter the plant roots and those that do, cannot establish a feeding site. To understand its ability to prevent RKNs from forming feeding sites, we performed transcriptomic analysis of *S. sisymbriifolium* roots inoculated with the Northern root knot nematode, *M. hapla*. Combined with the annotation of the recently published *S. sisymbriifolium* genome assembly, we discovered 13 differentially expressed resistance-related genes upon nematode inoculation. By transforming potatoes with candidate resistance genes from *S. sisymbriifolium*, we aim to understand the strong genetic resistance in *S. sisymbriifolium* and whether those genes are necessary and sufficient to drive resistance to RKN in potatoes. This information will help us understand gene functions and help us generate RKN resistance in relevant *Solanum* crops.

## Introduction

Plant parasitic nematodes (PPNs) are devastating pathogens of many crops, accounting for over $80 billion in annual yield losses in global agriculture (Nicol *et al*., 2011). Among PPNs, the sedentary endoparasites in the Tylenchida are well known for their agricultural impacts. Within this order, one of the most agronomically important groups of sedentary endoparasites is the root-knot nematode (RKN, *Meloidogyne* spp.). RKNs have a broad host range and a worldwide distribution (Kaloshian and Teixeira 2019); moreover, RKN’s unique life cycle has made it difficult to eradicate or even control. RKNs hatch as second stage juveniles (J2) from eggs and actively look for the host roots. After successfully penetrating the root, J2s migrate towards the root tip, make a U-turn, and enter the vascular cylinder to establish their feeding sites. By manipulating acytokinetic mitosis, the RKN J2s transform 5-12 vascular parenchyma cells into enlarged feeding sites called “giant cells” (Matuszkiewicz and Sobczak 2024). The nematode feeds on the contents of these giant cells using its needle-like mouth part called a stylet. Interestingly, RKN not only uses its stylet to feed from the giant cells but also to secrete molecules called effectors, many of which suppress the plant immune responses (Vieira and Gleason, 2019, Jagdale *et al.,* 2021, Bali and Gleason, 2024). Once they obtain enough nutrients, the J2s will halt feeding and molt three times to the young adult stage. Only females reestablish feeding where they continue to grow and produce eggs while males migrate out from the root. The plant cells around the female divide to form a “root gall” (Palomares-Rius *et al.,* 2017).

*Meloidogyne hapla* and *M. chitwoodi* are the predominant RKNs in the Pacific Northwest (PNW) region of the United States (Elling, 2013, Lima *et al.,* 2018, Zasada *et al.,* 2019). *Meloidogyne chitwoodi* and *M. hapla* have been found, respectively, in 22% and 14% of samples sent to local diagnostic clinics in the PNW (Zasada *et al*., 2018). Both pests can cause potato tuber blemishes, potentially making the tubers unmarketable (Zasada *et al*., 2019).

The most effective means to control RKNs are with chemical pesticides. Although nematicides/fumigants can be extremely effective in RKN control, there are increasing concerns about mammalian toxicity and environmental health (Bernard *et al.,* 2017). There are stringent restrictions on pesticide applications in agriculture, emphasizing the importance of environmental safety as well as human and animal health worldwide (Sasanelli *et al*., 2021; Desaeger *et al.,* 2020). Ideally, nematode management practices should include measures that are both environmentally friendly and economically sustainable. Genetic resistance is an environmentally friendly approach to controlling nematodes, but there is a limited number of known nematode resistance (*R*) genes and very few nematode *R* genes have been cloned (Rutter *et al.,* 2022).

In terms of potatoes, there is no genetic resistance to RKN in commercially available potato cultivars, and nematicides for nematode control are expensive for growers (Talavera-Rubia *et al.,* 2022). Considering that the thriving potato industry in the PNW has a net worth $1.8 BN (reported in USDA potato 2023 summary^1^), RKNs directly threaten the profits of the potato industry. Given the gravity of the situation, there is a critical need for diverse, effective, and sustainable management tools for RKNs.

One source of resistant material is the wild relatives of cultivated crops, which are usually well adapted to changing environments and pathogen attack, making them a rich source of pathogen resistance genes (Bradeen, 2021). For example, *Mi1-2* is an *R* gene that confers resistance in tomato to three species of RKN (*M. incognita*, *M. javanica*, and *M. arenaria*) (Milligan *et al.,* 1998). The resistance was discovered in the wild Solanum species *Solanum peruvianum* and introduced into cultivated tomatoes (Smith, 1944). *Mi-1.2* encodes a protein with coiled-coiled (CC), nucleotide binding (NB) and leucine-rich repeat (LRR) motifs (Milligan *et al.,* 1998). The Mi-mediated resistance is characterized by a hypersensitive response (HR) upon nematode infection, and the HR prevents nematode feeding site formation (Sato *et al.,* 2019).

Recent reports showed that *S. sisymbriifolium*, a wild Solanum plant, exhibited resistance to several *Meloidogyne* spp. (including *M. chitwoodi* and *M. hapla)* as well as to the cyst nematode (CN), *Globodera pallida* (Dandurand *et al.,* 2019, Hajihassani *et al.,* 2020, Perpétuo *et al.,* 2021, Baker *et al.,* 2023). *Solanum sisymbriifolium*, also called litchi tomato or sticky nightshade, is usually used as a PPNs trap crop to reduce PPNs population in the US (APHIS 2022^2^). For instance, *S. sisymbriifolium* stimulates *G. pallida* egg hatching, but these nematodes cannot reproduce in *S. sisymbriifolium* (Timmermans *et al.,* 2006). During parasitism, CNs inside the *S. sisymbriifolium* roots induce genes capable of promoting a HR (Kud et al 2021). This immune response leads to the cessation of feeding site formation, thereby preventing CN from completing their life cycle (Baker *et al.,* 2023). The cell death response shortly after feeding site initiation suggests a gene-for-gene response triggered by an *R* gene. Work with the two RKNs *M. chitwoodi* and *M. hapla* shows that *M. chitwoodi* can never enter the roots. Meanwhile, *M. hapla* rarely enters the roots and the juveniles that penetrate the roots never feed (Baker et al 2023). We hypothesize that the *M. hapla* J2 in the roots are triggering *R*-gene mediated cell death which prevents nematode feeding. Our long term goal is to identify the *R* gene(s) that are involved in *S. sisymbriifolium* resistance to *M. hapla.* To achieve this goal, we have developed genomic tools for *S. sisymbriifolium*. By utilizing transcriptomics and comparative genomic methods, we have laid the groundwork for *R* gene identification in *S. sisymbriifolium*.

## Materials and Methods

All common chemicals and reagents were purchased from Sigma Aldrich unless specified.

### Plant Material and nematode inoculation

Tissue culture of the *S. sisymbriifolium* clone SisSynII-1have been described previously (Casavant *et al*., 2017). The RKN susceptible potato cultivar used was *Solanum tuberosum cv.* ’Russet Burbank’. Plants were subcultured in modified Murashige and Skoog (MS) salts medium (pH 5.6, 0.43% (g/L) MS base salt (Sigma Aldrich M5519), 3% (g/L) sucrose, 0.7% (g/L) agar, 100 µg/mL myo-inositol, 2 µg/mL glycine, 1 µg/mL thiamine HCl, 0.5 µg/mL pyridoxine, and 0.5 µg/mL nicotinic acid) (Wixom *et al*., 2020) in a growth chamber at 22 °C with a 16 hour day light cycle. After three weeks of growth, the regenerated roots from the *S. sisymbriifolium* plants were cut off and replanted into a half MS salts medium (2.2% (g/L) MS base salt, 10% (g/L) sucrose, 0.05% MSV1, 0.7% (g/L) agar) to develop enough roots for nematode infection assays.

*Meloidogyne hapla* (VW9) was maintained on the *Solanum lycopersicum* cv. Rutgers under greenhouse conditions at Washington State University Greenhouse, Pullman, Washington, USA. To extract *M. hapla* eggs, the bulk root of *M. hapla* infected tomato was collected and cleaned in 0.6% sodium hypochlorite with 5 minutes of vigorous shaking. The solution was then passed through 45 mm (to filter larger particles) and 25 mm sieves (to capture the nematode eggs). To further purify these eggs, the material on the 25 mm sieves was collected into a 50mL tube filled with 45mL of 70% (g/L) sucrose solution and 5mL DI water. After centrifuging at 1500 rpm for 10 minutes, the egg solution mixture was poured onto a 25 µm sieve and the eggs were rinsed with DI water completely before collection. To prepare the sterile nematodes juveniles for potato and *S. sisymbriifolium* tissue culture inoculation, *M. hapla* eggs were surface sterilized three times with 0.6% sodium hypochlorite and rinsed liberally with DI water. Afterwards, eggs were placed in a modified Baermann funnel apparatus with hatching solution (0.1% Plant Preservative Mixture (PPM), 0.01% DDT in sterile DI water) (Zhang and Gleason, 2021). After 4 days of hatching, the sterile *M. hapla* J2 were collected. For inoculation in *S. sisymbriifolium* tissue culture for RNA-seq, approximately 800 *M. hapla* J2s were evenly distributed on the surface of the tissue culture media. For the acid fuchsin staining, approximately 300 *M. hapla* J2s were inoculated in *S. sisymbriifolium* tissue culture roots and infected roots were collected at 3- and 6-days post inoculation (dpi).

### Nematode acid fuchsin staining

The *M. hapla* infected roots were collected at 3- and 6-days post inoculation (dpi) and treated with 1.2% sodium hypochlorite for 5 minutes. The roots were then rinsed with running water for 5 minutes, stained in acid fuchsin solution (0.35% acid fuchsin, 25% acetic acid) and briefly boiled using a microwave. After cooling down, the root solution was removed, and the root was rinsed with water before adding 25% acetic acid solution and incubating for at least 20 minutes before visualization under a Zeiss AXI0 Observer.A1 Microscope.

### *Solanum sisymbriifolium* transcriptome sample collection

To obtain RNA-seq samples, *S. sisymbriifolium* root tissues at 3 and 6 dpi were collected (3dpi_infected, 3day_control, 6dpi_infected, and 6day_control). Roots from 4 to 5 plants were binned into one RNA-seq sample and we submitted three biological replicates for each condition. Total RNA was extracted using Total RNA Purification Maxi Kit (Norgen, #26800) following the manufacture’s protocol with an optional on-column DNA removal using Norgen’s RNase-free DNase I kit (#25710). The resulting RNAs were further concentrated and purified by using the RNA Clean-UP and Concentration Kit (Norgen, #23600). cDNA libraries were prepared by reverse transcribed total RNA using a ProtoScript® II First Strand cDNA Synthesis Kit (NEB, #E6560) with d(T)23 VN oligo and sent to Novogene for sequencing using Illumina Novaseq 6000 (PE 150bp with 6Gb read depth coverage).

### Structural and functional annotation of *S. sisymbriifolium* genome assembly

The genome annotation was conducted by a high-performance computer, “Kamiak” at Washington State University and a pre-built genome structural and functional annotation workflow^3^ (Zhang *et al.,* 2024) was adapted in this study. In brief, the *S. sisymbriifolium* genome assembly was obtained from the National Genomics Data Center (NGDC)^4^ database under accession ID “PI 487464 01” (NGDC Bioproject:PRJCA010759) (Wu *et al.,* 2023). The data was uploaded to the GenSAS (Humann *et al.,* 2019) server^5^ and the RepeatModeler tool was used to predict the repeat regions. The genome assembly was soft-masked using Bedtools (Quinlan and Hall, 2010). Genome quality was assessed by assembly BUSCO (Manni *et al.,* 2021) using the eudicot_odb10 dataset. Repetitive elements of the *S. sisymbriifolium* genome assembly were annotated using EDTA v2.0.0, (Ou *et al.,* 2019). GEMmaker v2.1.0 (Quinlan and Hall, 2010; Hadish *et al.,* 2022) was used to map the available *S. sisymbriifolium* RNAseq data from our study and Wixom *et al.,* (2018, 2020) to the soft-masked genome. The generated BAM files were used as HINT files for BRAKER1 (Hoff et al., 2016), an RNA-seq based genome structural annotator. Following recent suggestions (Gabriel et al., 2021), the annotation accuracy can be enhanced by running the protein homology-based genome annotator BRAKER2 (Brůna et al., 2021). Therefore, Viridiplantae OrthoDB (Kuznetsov et al., 2023), *S. tuberosum* v6.1, (Pham et al., 2020), and *S. lycopersicum* ITAG5.0 (Zhou et al., 2022) protein datasets (obtained from Phytozome^6^(Goodstein et al., 2012) were used as hints for BRAKER2 to annotate the genes. To utilize long reads RNA-seq data from Wixom *et al.,* (2018), the long read integration protocol BRAKER^7^ was used. Later, TSEBRA (Gabriel et al., 2021) was used to combine the BRAKER1, BRAKER2, and long-reads results. Lastly, structural annotated *S. sisymbriifolium* genome assembly was used for EnTAPnf (Hart et al., 2020) for *R* gene functional annotation with gene ontology assignment. The completeness of the functionally annotated genome assembly was assessed using BUSCO (Manni et al., 2021) referencing against the solanum_odb10 database and eudicot_odb10 dataset.

### Genome scaffolding and Synteny analysis

Due to the current absence of chromosome scaffolding information for the *S. sisymbriifolium* contig-level genome assembly, we used *S. wrightii* (NGDC Bioproject:<u>PRJCA010759</u>) as the reference chromosome genome and built a pseudochromosome-level genome assembly for *S. sisymbriifolium* using RagTag v2.1.0 (Alonge *et al*., 2022). Synteny analysis was performed using GENESPACE v.1.2.3 (Lovell *et al*., 2022) comparing a *S. sisymbriifolium* pseudochromosome-level genome with *S. tuberosum* v6.1 (Pham *et al*., 2020), *S. lycopersicum* ITAG5.0 (Zhou *et al*., 2022), *S. wrightii* (NGDC Bioproject:<u>PRJCA010759</u>), and *Solanum torvum* (NGDC Bioproject: <u>PRJCA010759</u>) (Wu *et al*., 2023).

### Differential gene expression and gene enrichment analysis

GEMmaker v2.1.0 (Quinlan and Hall, 2010; Hadish *et al*., 2022) was used to trim and map the *S. sisymbriifolium* RNAseq to the soft-masked genome. The resulting GEM files were processed with DEseq2 tool (Love, Huber and Anders, 2014) to identify differentially expressed genes (DEGs). DEGs with Log2 Fold change > 0.5 or <-0.5 with adjusted p-value <0.05 were selected for analysis. Gene ontology and Intropro scan term enrichment analysis was performed using func-e (‘SystemsGenetics/FUNC-E: Release v2.0.1’, no date).

### Real time quantitative PCR validation

To verify the DEGs, qRT-PCR was performed using the primer sets listed in Supplemental Table S1. An actin gene (Casavant *et al*., 2017) and elongation factor 1 Alpha (EF-1a) gene were selected as the reference genes in *S. sisymbriifolium* as both genes were reported as ubiquitous housekeeping genes (Stürzenbaum and Kille, 2001). PowerTrack™ SYBR Green Master Mix (Applied Biosystems™ #A46113) was used for the reactions. The qRT-PCR conditions are, 95°C for 3 min, 40 cycles of 95°C for 15 sec, 52°C for 15 sec and 72°C for 20 sec and followed by a melting curve analysis from 65°C to 95°C with 0.5°C increased incrementally at 5 seconds increments. The qRT-PCR validation was conducted twice using the cDNA samples submitted for RNA-seq and the cDNA obtained from another independent experiment. Each qRT-PCR consisted of three technical replicates for each of three bio replicates. The 2–ΔΔCt method (Livak and Schmittgen, 2001) was used to calculate the relative expression levels of each gene and Wilcoxon rank sum exact test was performed to examine the relative expression difference.

## Results

### *Solanum sisymbriifolium* can be infected by low numbers of *M. hapla* at 3 dpi

Transcriptome analysis of CN-infected *S. sisymbriifolium* was performed at 3 dpi (Wixom et al 2018). Baker et al (2023) previously showed that there were few *M. hapla* J2 inside the *S. sisymbriifolium* roots at 7 dpi. Earlier infection timepoints had not been assessed with *M. hapla*. Thus, we measured the level of *M. hapla* infections of *S. sisymbriifolium* and a control susceptible plant (potato) at an early timepoint (3 dpi) and a later time point (6 dpi). We inoculated the *S. sisymbriifolium* (Fig.1A) and RKN-susceptible potato Russet Burbank (Fig.1B) with 300 *M. hapla* J2s (6 bio-reps). At 3 and 6 dpi, significantly more J2s were found in potato (average 4.33 J2 per plant at 3dpi and 11.86 J2 at 6dpi) compared to *S. sisymbriifolium* (average 1.17 J2 per plant at 3dpi and 3.43 J2 at 6 dpi) (Fig.1C). This confirmed that the *S. sisymbriifolium* can be infected by a small number of *M. hapla* at 3 dpi. As a result, we subsequently performed the transcriptome analysis on *M. hapla* infected-*S. sisymbriifolium* at 3 dpi, similar to the previously performed transcriptome analysis of CN-infected *S. sisymbriifolium*.

**Figure 1.**
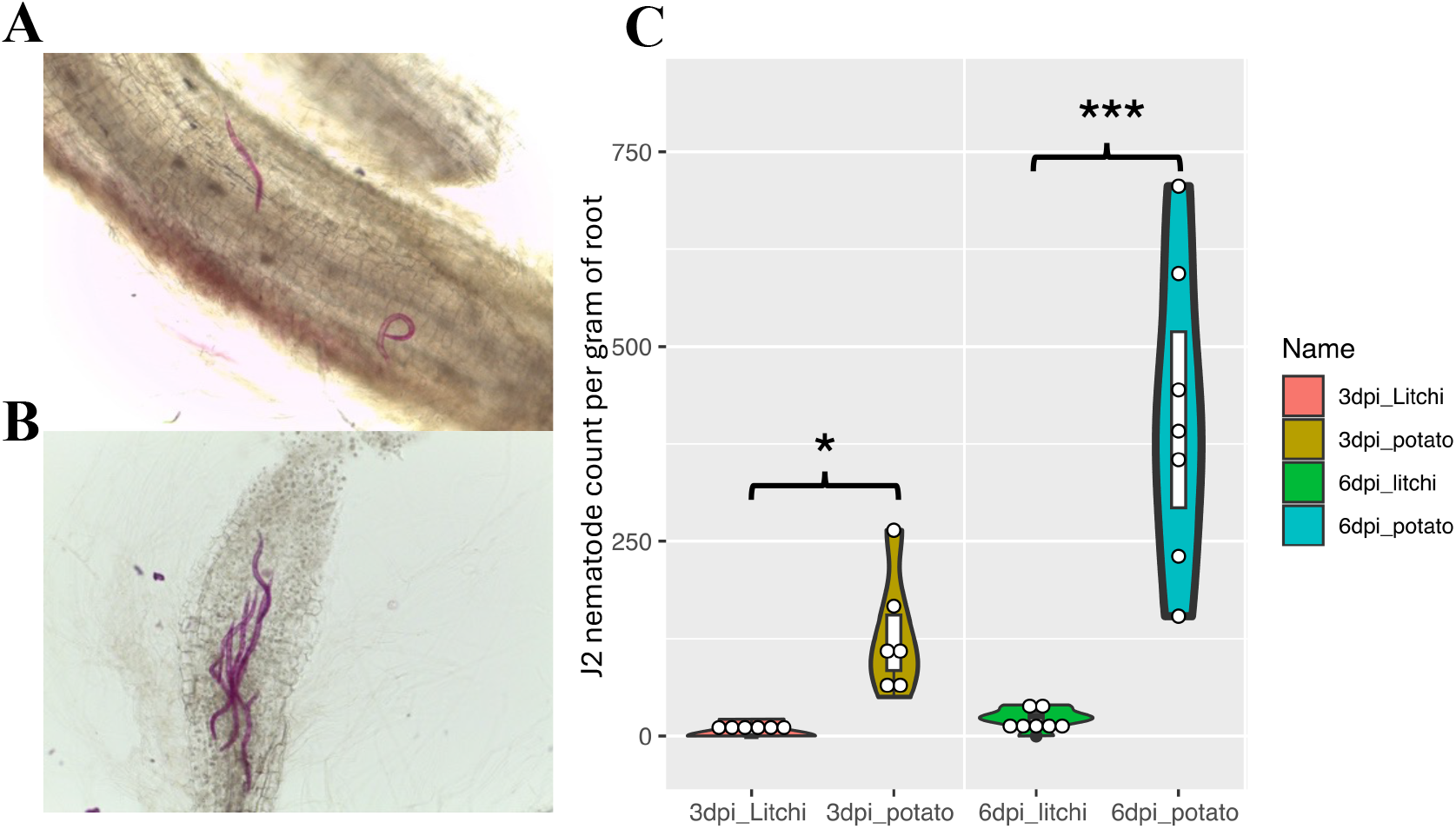
A. Acid Fuchsin staining of *S. sisymbriifolium* (A) and potato (B) roots at 3 dpi. The *M. hapla* J2s stain red. 100X magnification. C. Number of *M. hapla* J2 present in RKN-resistant *S. sisymbriifolium* (litchi) as compared to RKN-susceptible potato Burbank (potato) at 3 dpi and 6 dpi. Graphs show average number of J2 per gram of root +/- standard deviation. One way ANOVA test was performed for significant difference (*: p < 0.05, ***: p<0.001)

### Improved *S. sisymbriifolium* genome annotation combining existing de novo transcriptome and genome assembly. and transcriptome analysis

#### Genome reannotation

Recently, a contig level genome assembly of *S. sisymbriifolium* was published (Wu *et al*., 2023) with a 98.1% assembly BUSCO score, compared to the 88% assembly BUSCO score from the SMRT de novo transcriptome (Wixom *et al*., 2018) (Supplemental table S2),. To enhance the chance to identify possible RKN resistance genes in *S. sisymbriifolium*, we structurally and functionally annotated the *S. sisymbriifolium* genome assembly. *S. sisymbriifolium* genome assembly contains 78.47% of repeated sequences (Supplemental table S3), compared to 66.8% in potato (Pham *et al*., 2020) and 62.1% in tomato (Zhou *et al*., 2022).

To refine the gene model annotation, we utilized homolog protein datasets from closely related species and RNA-seq data generated in the Wixom *et al., (*2018) study and in this study using BRAKER tool as evidence for gene structure predictions (Fig. 2). To further improve the annotation results, we utilized the iso-seq long read RNA-seq data obtained from Wixom et al. (2018) (PRJNA422280), as extra data for BREAKER to predict accurate gene structure. In total, we identified 62,664 genes from the *S. sisymbriifolium* genome assembly (Wu *et al*., 2023) (Supplemental table S4), compared to 50,940 genes previously identified in *S. sisymbriifolium* SMRT transcriptome (Wixom *et al.,* 2018), 32,917 genes in the potato v6.1 genome (Pham *et al*., 2020), and 36,648 genes in the tomato ITAG5.0 genome (Zhou et al., 2022). There were 53,418 genes (85.2%) of *S. sisymbriifolium* genome assigned with at least one functional term (GO term, KEGG pathway, InterPro term, EggNOG orthogroup, NCBI refseq homolog). To assess the completeness of this genome annotation, the completed BUSCO annotation score for the final annotation data is 95.8 % using odbSolanium 10 database (95.1% using odbEudicot10 database), which is better than current tomato and potato reference genomes BUSCO annotation scores (Table 1).

**Figure 2.**
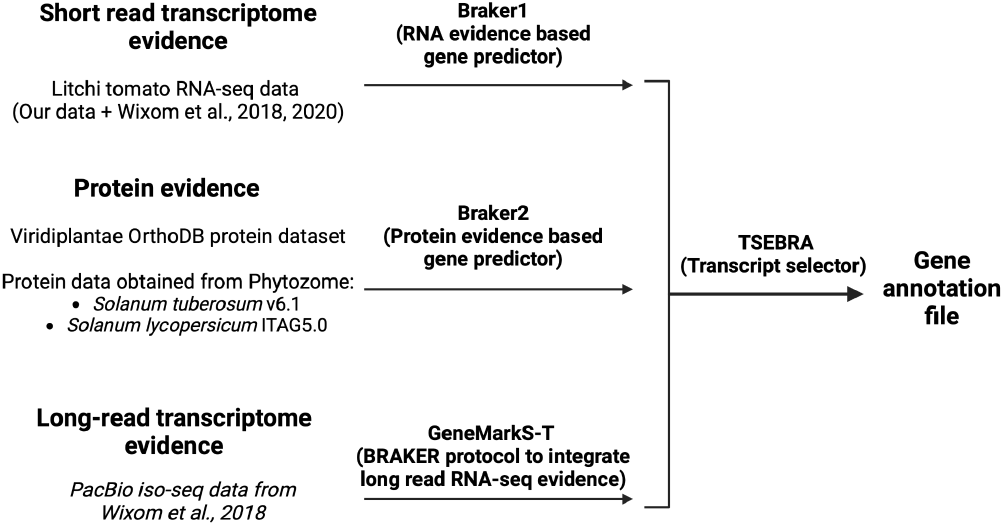
Schematic chart of genome annotation method. Figure created with BioRender.com

**Table 1.** Summary of annotation BUSCO scores of *S. sisymbriifolium*, potato, and tomato genomes.

| Input_file | Dataset | Complete | Single | Duplicated | Fragmented | Missing | n_markers |
| --- | --- | --- | --- | --- | --- | --- | --- |
| Slycopersicum_796_ITAG5.0.protein.fa | solanales_odb10 | 94.8 | 80.4 | 14.4 | 0.9 | 4.3 | 5950 |
| Stuberosum_686_v6.1.protein.fa | solanales_odb10 | 95.5 | 68.8 | 26.7 | 1.9 | 2.6 | 5950 |
| Slycopersicum_390_ITAG2.4.protein.fa | solanales_odb10 | 94.6 | 92.6 | 2 | 2.4 | 3 | 5950 |
| Stuberosum_448_v4.03.protein.fa | solanales_odb10 | 81.7 | 61.9 | 19.8 | 2.7 | 15.6 | 5950 |
| litchi_softmaskedv2.fa.protein.faa | solanales_odb10 | 95.8 | 85.1 | 10.7 | 0.7 | 3.5 | 5950 |

#### Synteny analysis with other relatives

Next, to understand the evolutionary relationship of *S. sisymbriifolium* and its close relatives, synteny analysis was performed. Due to the unavailability of a chromosome level genome assembly, *S. sisymbriifolium* pseudochromosomes were assembled based on the *S. wrightii* genome which is the most closely related species that has chromosome level assembly (Wu *et al*., 2023). The gene contents of *S. sisymbriifolium* pseudochromosomes and *S. wrightii* chromosome 6 and 8 were presented in potato and tomato chromosomes 4 and 11 due to a potential inter-chromosomal arrangement (Fig 3) during the Solanum genus speciation about 22-29 Million Years Ago (D’Agostino *et al*., 2013; Wu *et al*., 2023). When the potato genome was used as a reference to assemble the *S. sisymbriifolium* pseudochromosomes genome, most gene blocks aligned to chromosome 4 in potato and tomato, and multiple gene blocks in *S. sisymbriifolium* pseudochromosome 4 are syntenic to the regions of potato and tomato chromosome 10 and 11 (Supplemental Figure S1). This observation implies that an inter-chromosomal arrangement event in two chromosomes (chromosomes 4 and 11 in potato and tomato) had happened between the common ancestors of wild Solanum species (*S. sisymbriifolium, S. wrightii, and S. torvum)* and that of cultivated Solanum species (potato and tomato).

**Figure 3.**
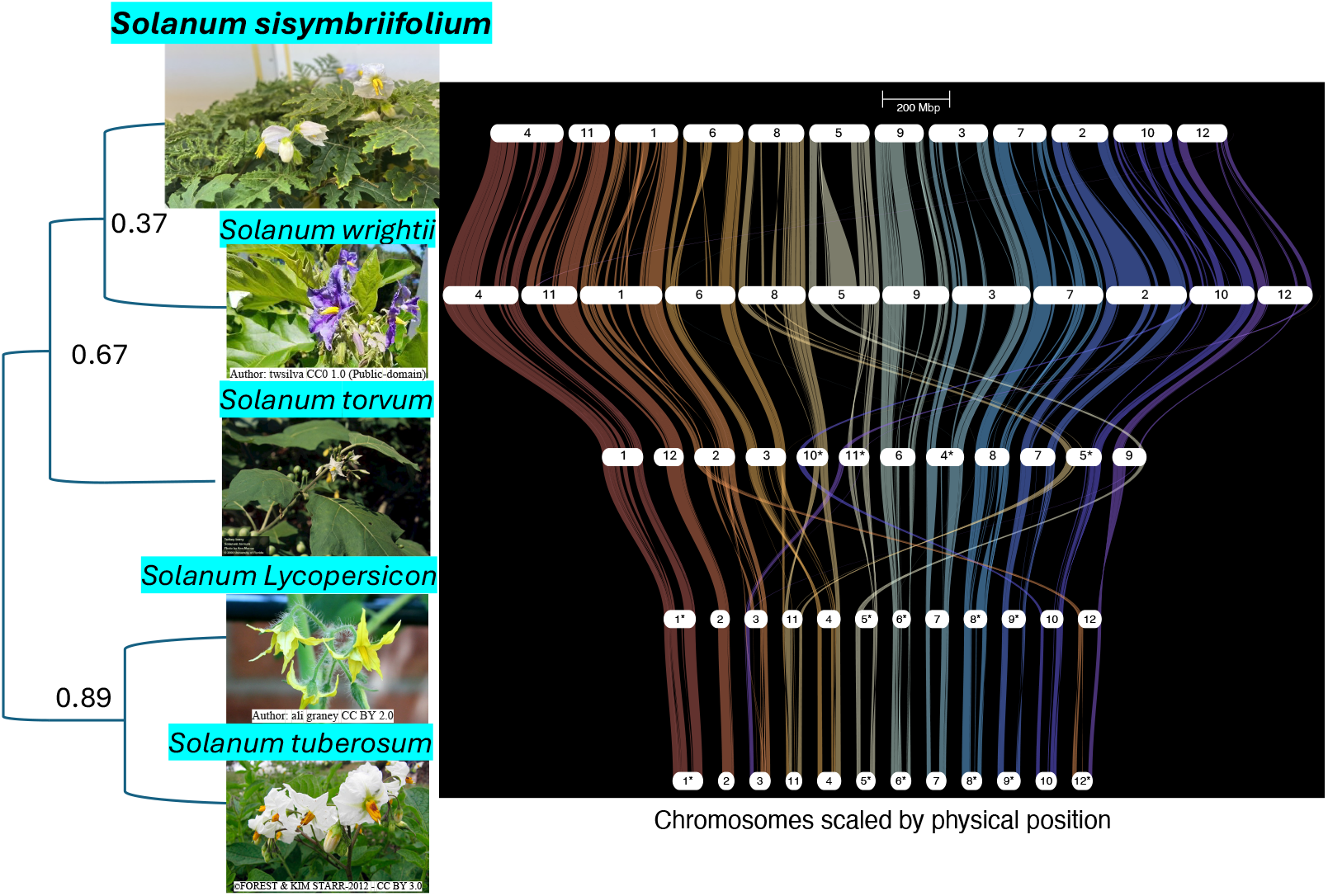
Synteny analysis of *S. sisymbriifolium* pseudochromosomal genome based on the *S. wrightii* reference genome, compared to the other Solanum species, scaled by chromosome physical location. Left: Species tree based on OrthoFinder result (Emms and Kelly, 2019). Numbers on the left represent bootstrap relatedness.

#### Transcriptome analysis of *S. sisymbriifolium* inoculated with *M. hapla*

To understand the *S. sisymbriifolium* gene response to RKN infection, we analyzed the RNA-seq data of *S. sisymbriifolium* infected with *M. hapla* and a mock control for differentially expressed genes (DEGs). More DEGs were identified within the 3 dpi dataset (843 genes) compared to the 6 dpi dataset (230 genes) with only a few overlapping genes between the timepoints (Supplemental Table S5, S6). In the 3dpi dataset, the most enriched term in upregulated genes is protein phosphorylation (GO:0006468), while downregulated genes are enriched for the obsolete oxidation-reduction process (GO:0055114). Similarly, in the 6 dpi dataset, the most enriched term for upregulated genes includes the regulation of photoperiodism, flowering (GO:2000028), while downregulated genes are enriched for the obsolete oxidation-reduction process (GO:0055114). A full list of GO terms and Interpro Scan terms enrichment is available in Supplemental Table S7

Notably, UMAP analysis segregates the RKN infected samples from the control group within the 3 dpi dataset, as opposed to the 6 dpi dataset (Fig 4). Collectively, we hypothesize that *S. sisymbriifolium* responses to *M. hapla* infection were more active at the early time point (3 dpi).

**Figure 4.**
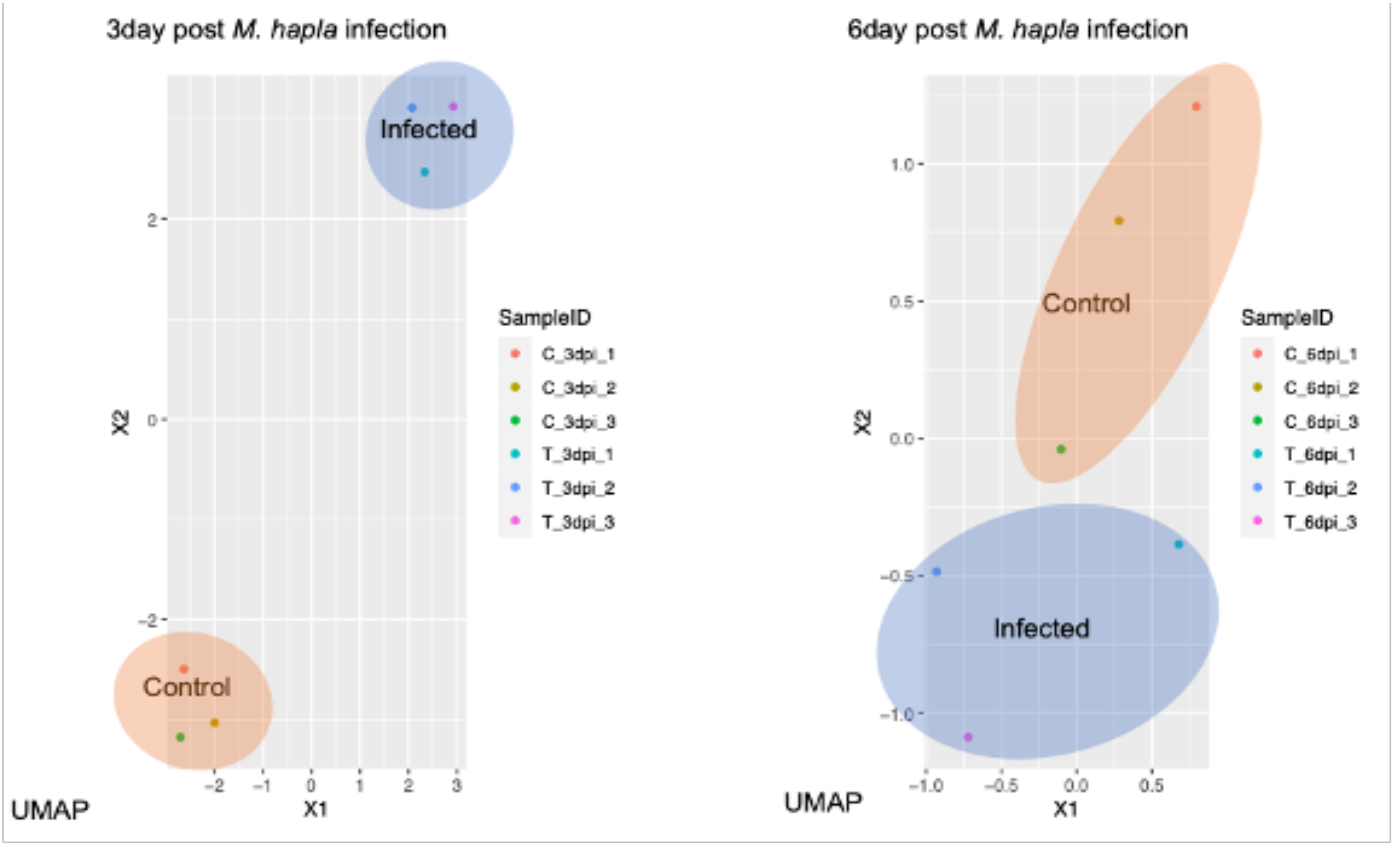
The UMAP plot reveals a distinct clustering pattern within the 3dpi dataset, where all the RKN infection and control bio-replicates are organized into separate clusters. In contrast, the distinction becomes less pronounced at 6 dpi.

To determine if *S. sisymbriifolium* responds differently to RKN in comparison to CN, we compared our *M. hapla* infected *S. sisymbriifolium* RNA-seq data with previously the published *G. pallida* infected *S. sisymbriifolium* RNA-seq data (Wixom *et al*., 2020) reanalyzed with the newly annotated *S. sisymbriifolium* genome. Ninety-four genes were upregulated in both transcriptomes of cyst and root-knot nematodes infected *S. sisymbriifolium* with the most enriched term being “oxidoreductase activity” (GO:0016491) and 164 genes were down regulated with the most enriched term being “membrane” (GO:0016020) (Fig.5). A full list of DEGs shared in the transcriptomes of cyst and root-knot nematodes infected *S. sisymbriifolium* and a Full list of their GO terms and Interpro terms enrichment is available in Supplemental Table S8 and S9.

**Figure 5.**
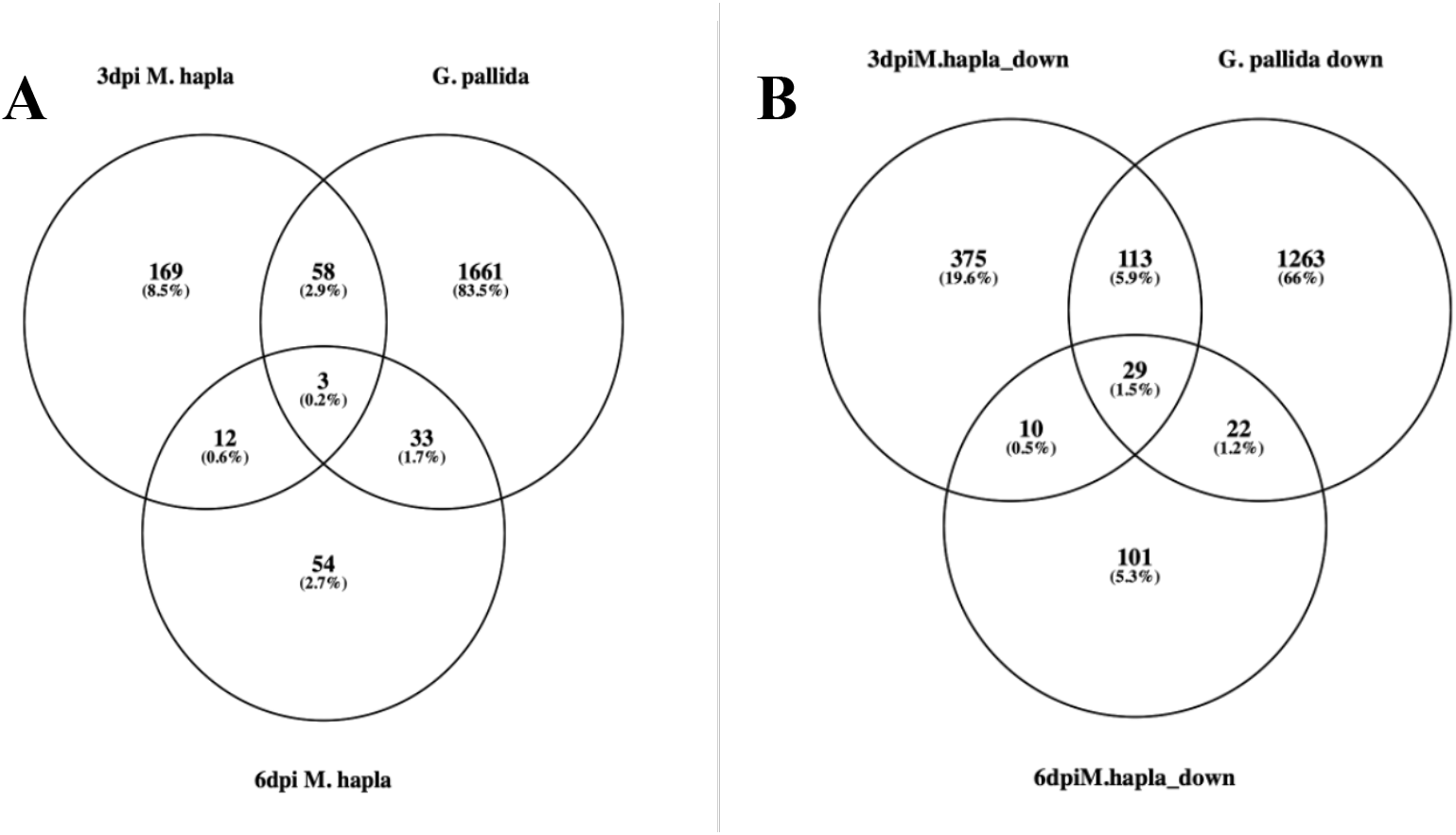
Overlap of upregulated (A) and downregulated (B) DEGs of *M. hapla* infected *S. sisymbriifolium* RNA-seq data (3dpi and 6dpi) with previously published *G. pallida* infected *S. sisymbriifolium* RNA-seq data (3dpi) (Wixom *et al*., 2020)

Transcriptome analysis of *S. sisymbriifolium* infected by the CN *G. pallida* at 3-dpi implied that Reactive Oxygen Species (ROS) and the ethylene production pathways are responsible for CN resistance (Wixom *et al*., 2020). From our data, many down regulated genes (especially at 3 dpi) were related to ROS scavengers (antioxidative genes e.g. peroxidase (APX), catalase (CAT), Thioredoxin/Peroxiredoxin Q (PrxQ) etc.), which might imply that *S. sisymbriifolium* maintained at high ROS level and may be related to the induction of plant defenses by the nematode. A similar inhibition of ROS scavenger gene expression was found in resistant *Mi1.2*-tomato upon *M. incognita* infection, and the inhibition of ROS-scavenging systems may contribute to the HR around the head of the nematode shortly after infection (Molinari and Leonetti 2023).

In the case of *S. sisymbriifolium* responses to *M. hapla* infection, we discovered a diverse group of putative nucleotide binding-leucine rich repeat (NB-LRR) genes that were significantly differentially expressed (Table 2). From the list, only Sol.si.v1a2.GWHBKCF00000002.g002740 was upregulated in *S. sisymbriifolium* upon both *M. hapla* (RKN) and *G. pallida* (CN) infection at 3 dpi.

**Table 2.**
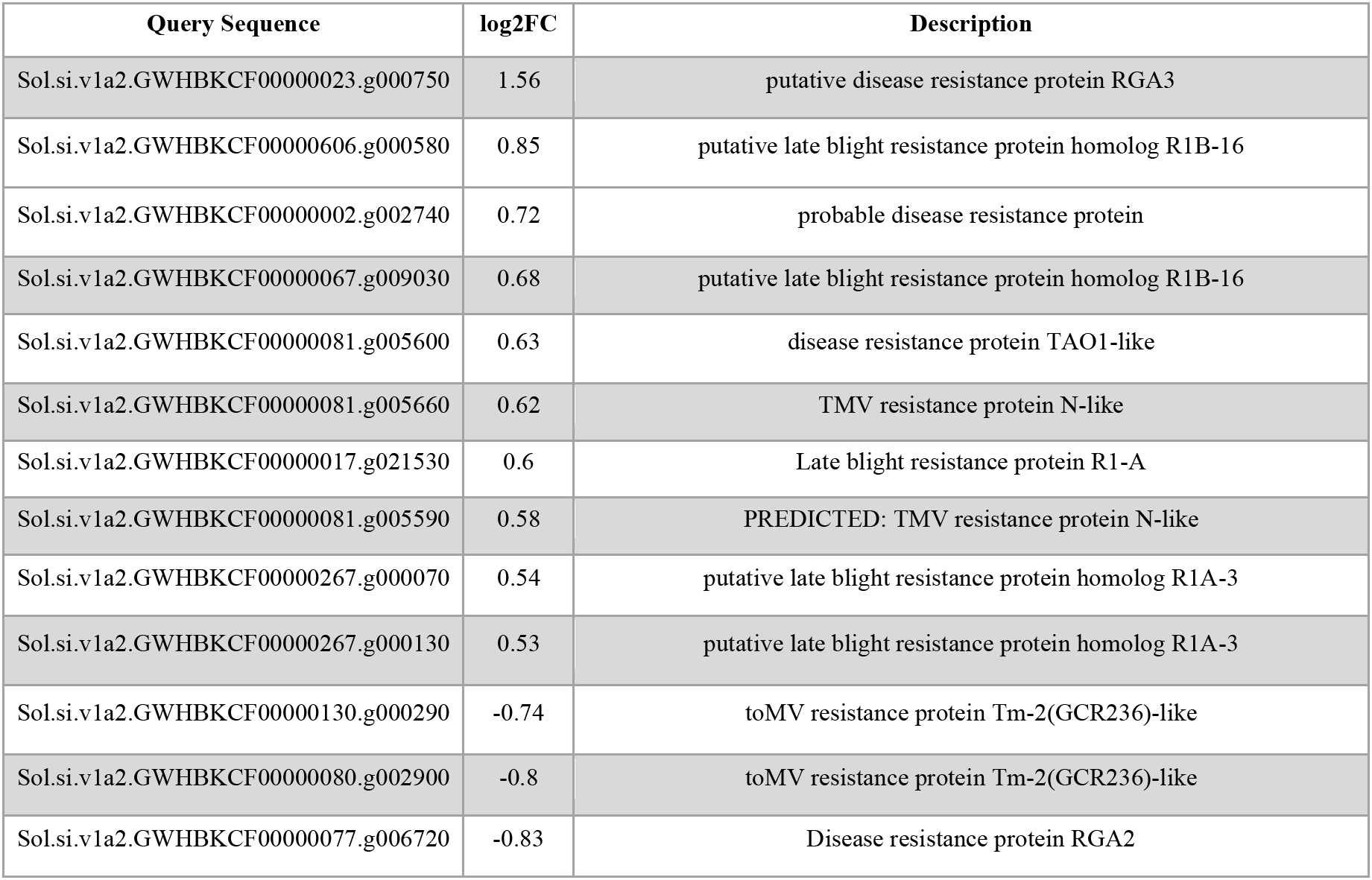
13 putative resistance-related genes were differentially expressed in *S. sisymbriifolium* infected by *M. hapla*.

To validate the upregulation of these putative *NB-LRR* genes upon nematode infection, we selected the top 4 genes for expression analysis by qRT-PCR: Sol.si.v1a2.GWHBKCF00000023.g000750, Sol.si.v1a2.GWHBKCF00000606.g000580, Sol.si.v1a2.GWHBKCF00000002.g002740, and Sol.si.v1a2.GWHBKCF00000067.g009030. We verified that all 4 genes were transcriptionally upregulated in litchi tomato upon *M. hapla* infection using cDNA derived from the RNA sent for RNA-seq (Fig.6 A, Supplemental Figure S2A). This confirmed the gene expression data from the RNAseq bioinformatics pipeline. However, from the transcriptome analysis, all 4 genes had relatively low, but significant values for fold change. We reasoned that since only a few nematodes could enter *S. sisymbriifolium* roots at 3 dpi (Fig 1), only a small proportion of the roots are responding to nematode infection. Therefore, using bulk root sample for RNA-seq could dilute the utility of the plant gene expression analysis. To eliminate potential artifacts from expression analysis on a single experiment, we performed an independent qRT-PCR experiment using cDNA synthesized from the RNA of a separate infection experiment. However, only Sol.si.v1a2.GWHBKCF00000606.g000580 and Sol.si.v1a2.GWHBKCF00000002.g002740 were upregulated (Fig.6 B, Supplemental Figure S2B). This confirms that these two genes are robustly up-regulated in expression during *M. hapla* infection and warrant further investigation.

**Figure 6.**
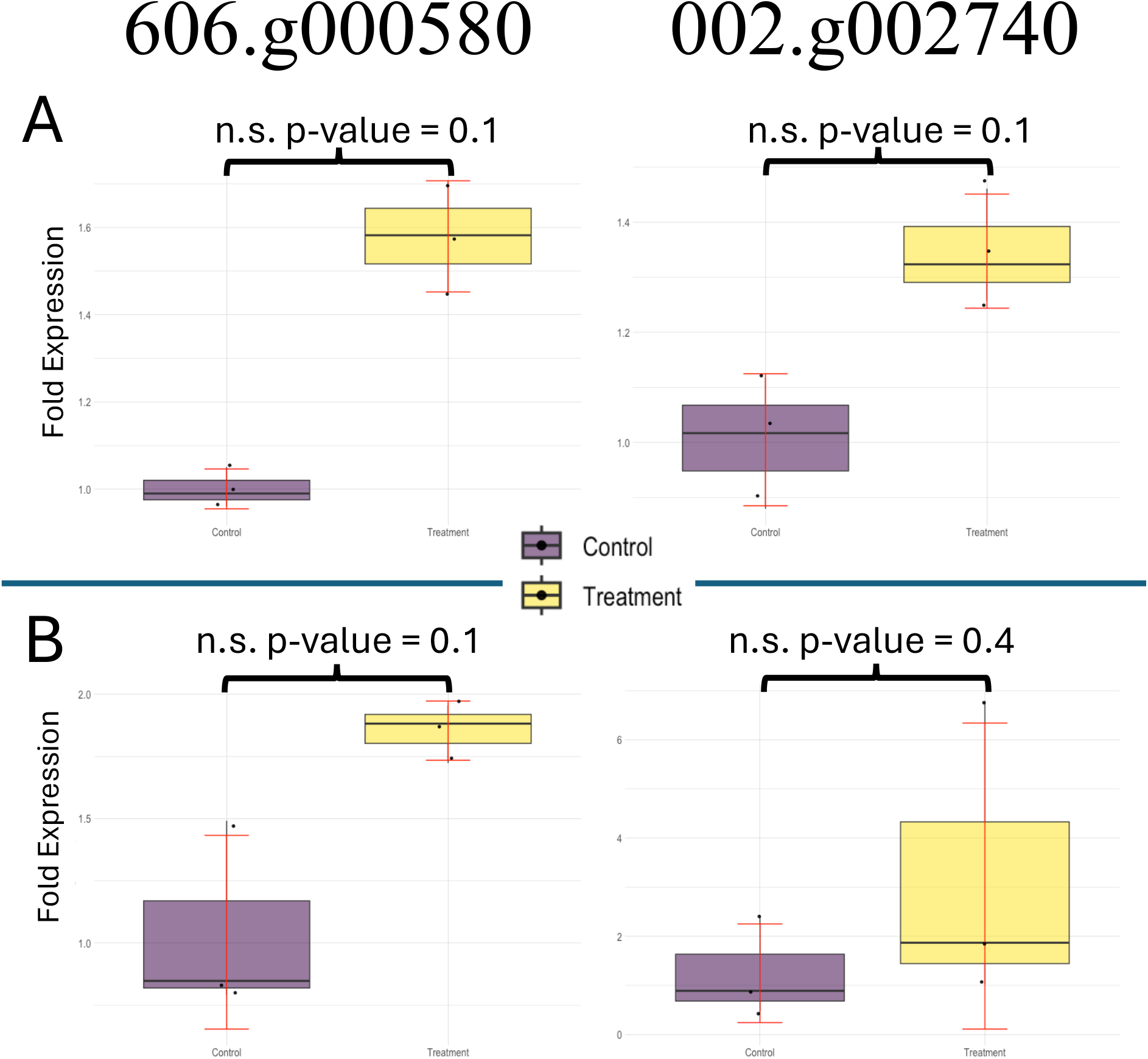
The expression level of 2 upregulated *S. sisymbriifolium* putative NB-LRR genes after *M. hapla* infection and mock condition: Sol.si.v1a2.GWHBKCF00000606.g000580 and Sol.si.v1a2.GWHBKCF00000002.g002740. The box plots show average fold change relative to the mock control. Error bar indicates standard deviation. Each back dot represented a single bio replicate (n =3). The experiment was done twice: A) qRT-PCR was performed using cDNA synthesized from the RNA used for RNA-seq. B) qRT-PCR was performed using cDNA synthesized from the RNA isolated from an independent *S. sisymbriifolium - M. hapla* infection experiment. Wilcoxon rank sum exact test was performed for significant difference. n.s.: no significant difference.

## Discussion

Overall*, Solanum sisymbriifolium* exhibits broad resistance against plant parasitic nematodes, including two RKN species, *M. chitwoodi* and *M. hapla*, which are significant pests in potato production within the Pacific Northwest (PNW). Given the absence of genetic resistance to RKN in cultivated potato varieties, exploration of wild potatoes and distantly related Solanum species presents a promising avenue for identifying effective *R* genes transferable to potato for enhanced nematode control.

Previously, Liu *et al*. (2015) discovered that *Solanum linnaeanum*, another wild relative of potatoes, contains an *R* gene that targets Verticillium wilt and successfully transferred its resistance into eggplant through hybridization. This intriguing result indicates that transferring an *R*-gene from wild Solanum plants into a distantly-related, cultivated Solanum crop can provide pathogen resistance to a previously susceptible plant (Liu, Zheng *et al.,* 2015).

The identification of potential R candidates from the highly resistant *S. sisymbriifolium* necessitated a comprehensive genome annotation and transcriptome analysis of both infected and uninfected roots. This allowed us to identify *R* gene candidates that are differentially expressed. Future work will include the functional characterization of selected NB-LRR proteins by their introduction into susceptible potato cultivars via Agrobacterium-mediated transformation.

Furthermore, leveraging the chromosomal synteny shared between potato and *S. sisymbriifolium* may offer a strategic approach to delineating syntenic chromosomal blocks harboring resistance genes. This comparative genomics approach may help facilitate the identification of homologous potato genes for further investigation and enhance our understanding of the genetic basis underlying resistance mechanisms.

## Acknowledgement

Thanks to Dr. Stephen Ficklin and Dr. Huiting Zhang for providing computation resources and technical advice for genome annotation. This research is funded by in part by the Northwest Potato Research Consortium and USDA National Institute of Food and Agriculture (NIFA), award number 2022-51181-38450.

## Author contributions

I.K. and C.G. contributed to the experiment design. I.K., A.C., J.K. and C.G. contributed to data analysis, and manuscript writing. I.K. conceived the experiments and data collection. All authors contributed to the article and approved the submitted version

## Data availability

The *S. sisymbriifolium* genome annotation, pseudo-scaffolding genome, and RNA-seq data have been deposited at the NCBI database^8^ under BioProject ID XXXXXXX.

## Supporting Information

Supplementary material is available at XXXXXX

---

**Supplemental figure S1.**
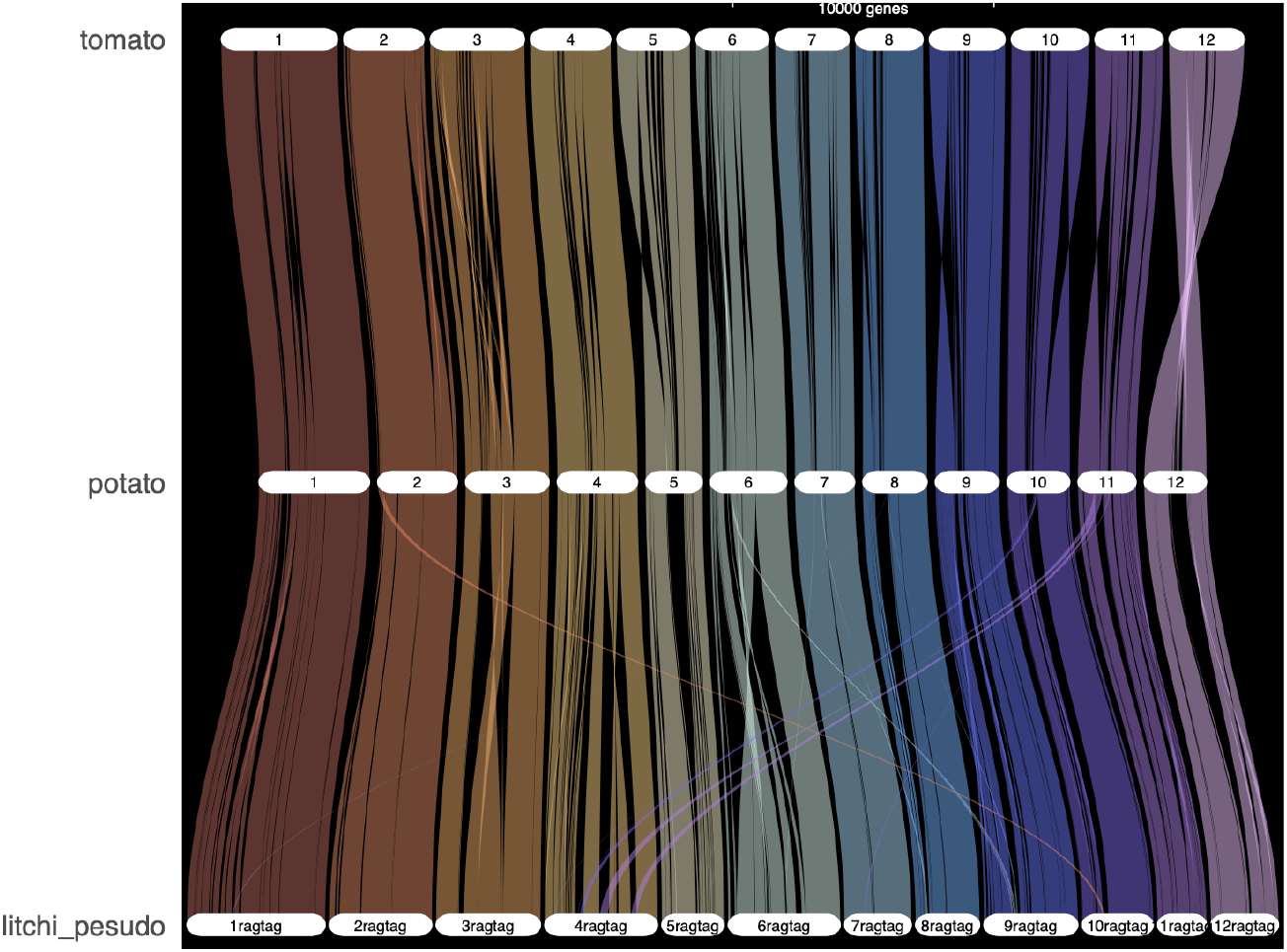
Synteny analysis of *S. sisymbriifolium* pseudochromosomal genome (litchi_pesudo) based on the potato reference genome, scaled by gene rank order.

**Supplemental Figure S2.**
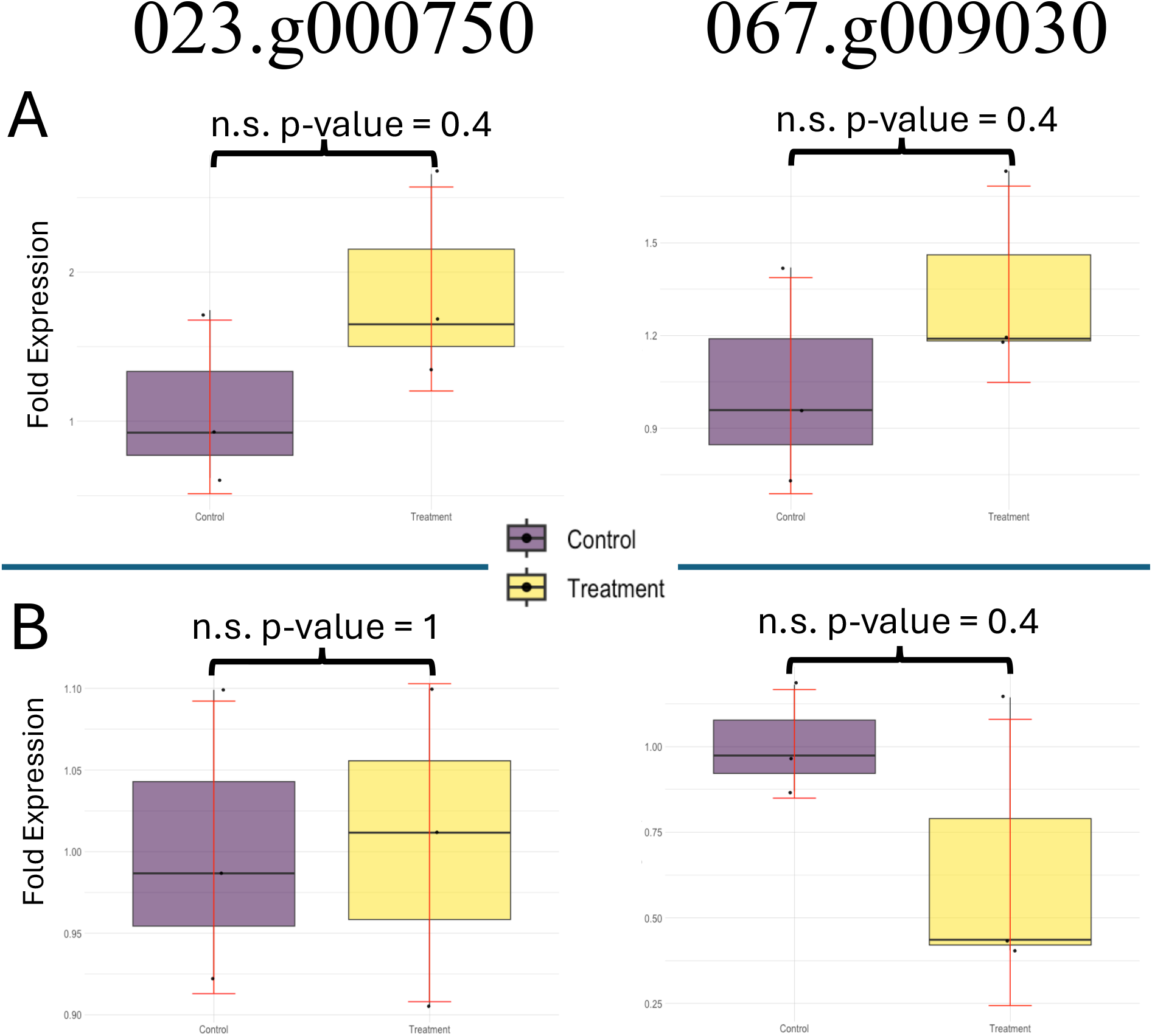
The expression level of 2 not upregulated S*. sisymbriifolium* putative NB-LR*R gene*s under *M. hapla* infection and mock condition: Sol.si.v1a2.GWHBKCF00000023.g000750 and Sol.si.v1a2.GWHBKCF00000067.g009030. The graphs show fold change relative to the mock control in a box plot with mean value. Error bar indicates standard deviation. Each back dot represented a single bio replicate (n =3). The experiment was done twice: A) First round qRT-PCR was performed using the total RNA samples used for RNA-seq. B) Total RNAs used for second round of qRT-PCR were obtained from a separate *S. sisymbriifolium* - M. hapla infection experiment. Wilcoxon rank sum exact test was performed for significant difference. n.s.: no significant difference.

## Footnotes

1 https://usda.library.cornell.edu/concern/publications/fx719m44h

2 https://www.aphis.usda.gov/aphis/newsroom/stakeholder-info/stakeholder-messages/plant-health-news/preliminary-litchi-tomato-trap-crop-weed-risk-analysis-webinar

3 https://gitlab.com/ficklinlab-public/wa-38-genome

4 https://ngdc.cncb.ac.cn

5 https://www.gensas.org/

6 https://phytozome-next.jgi.doe.gov/

7 https://github.com/Gaius-Augustus/BRAKER/blob/master/docs/long_reads/

8 https://www.ncbi.nlm.nih.gov/bioproject/

